# scDisent: regulatory-aware disentangled representation learning for multi-omic single-cell analysis

**DOI:** 10.64898/2026.04.12.717909

**Authors:** Guoren Xi

## Abstract

Single-cell multi-omic technologies measure complementary aspects of cellular identity and regulatory state, yet most integration models compress these signals into one entangled latent space. Such representations are useful for clustering but poorly suited to regulator-centered interpretation or perturbation-oriented analysis. We present scDisent (https://github.com/xiguoren/scDisent), a generative framework that separates expression-associated variables (***z***_***expr***_) from regulation-associated variables (***z***_***reg***_) and links them through a sparse directed mapping. scDisent combines modality-specific encoding, variational disentanglement, total-correlation and orthogonality regularization, and a Gumbel-gated causal module protected by detach-based gradient isolation. Across benchmark datasets with matched modalities, scDisent achieved the strongest clustering performance among the tested methods while exposing regulatory structure that competing integration models do not represent explicitly. The learned causal atlas remained sparse, perturbation analyses recovered biologically coherent lineage-associated programs, and branch-separation analyses showed that benchmark-label information concentrated in ***z***_***expr***_ rather than ***z***_***reg***_. These results position scDisent as a multi-omic representation model that improves both integration quality and biological interpretability.

## 1 Introduction

Single-cell multi-omic assays make it possible to observe complementary molecular layers within the same cell and have motivated a large class of integration methods for denoising, alignment and cell-state discovery (Clark et al. 2018; Chen et al. 2019; Ma et al. 2020; Swanson et al. 2021; Mimitou et al. 2021; Stuart et al. 2019; Stuart and Satija 2019; Argelaguet et al. 2020; Liu et al. 2020; Gong et al. 2021; Lin et al. 2022; Korsunsky et al. 2019; Zhu et al. 2020, 2023). These approaches are effective for clustering and manifold learning, but they usually optimize a single latent space that mixes identity-associated and regulation-associated variation.

This entanglement limits biological interpretation. If identity and putative regulatory drivers occupy the same coordinates, latent perturbation becomes difficult to interpret mechanistically. The issue spans variational, factor-analysis and graph-guided models, including scVI, MultiVI, MOFA+, LIGER, Cobolt, scJoint, Harmony, scGLUE and WNN (Lopez et al. 2018; Ashuach et al. 2023; Argelaguet et al. 2020; Liu et al. 2020; Gong et al. 2021; Lin et al. 2022; Korsunsky et al. 2019; Cao and Gao 2022; Hao et al. 2021). Translation-oriented models such as BABEL and Polar-bear support cross-modal prediction, but they do not provide an explicit split between identity-preserving and regulation-oriented latent factors (Wu et al. 2021; Zhang et al. 2022).

We address this problem with scDisent, a disentangled representation model with directed regulatory mapping for single-cell multi-omic analysis. The model factorizes the latent space into an expression branch, *z*_*expr*_, and a regulatory branch, *z*_*reg*_, and links them through a sparse directed mapping. This design aims to preserve an identity anchor while exposing a second branch better suited to regulator-centered interpretation and in silico perturbation (Chen et al. 2018; Argelaguet et al. 2021; Zhu et al. 2023).

Across PBMC, Human Brain and Mouse E18 benchmark datasets with matched modalities, scDisent achieved the strongest clustering performance among the tested methods while producing sparse directed regulatory mapping, interpretable perturbation outputs and quantitative evidence of branch separation. The study therefore asks not only whether integration improves, but whether the learned representation becomes more useful for biological reasoning.

## 2 Methods

### 2.1 Problem formulation

Let *X*_*rna*_ *∈ ℝ*^*n×p*^ and *X*_*atac*_ *∈ ℝ*^*n×q*^ denote RNA and ATAC views of the same cells. Rather than learning one mixed embedding, scDisent factorizes the latent state as

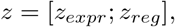

where *z*_*expr*_ preserves expression-associated identity and *z*_*reg*_ captures regulation-oriented variation. The model remains a multimodal variational autoencoder,

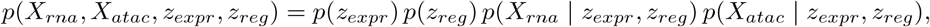

with an additional sparse mapping from *z*_*reg*_ to *z*_*expr*_ that acts as a representation-level inductive bias rather than a claim of direct causal identification (Argelaguet et al. 2021; Zhu et al. 2023).

### 2.2 Datasets and baselines

We evaluated scDisent on PBMC 10k, Human Brain 3k, and Mouse E18 (10x Genomics 2020b, a, 2021). These benchmarks span immune, neural and developmental contexts and use internally generated reference benchmark labels derived during preprocessing (Supplementary Methods S1).

**Table 1.**
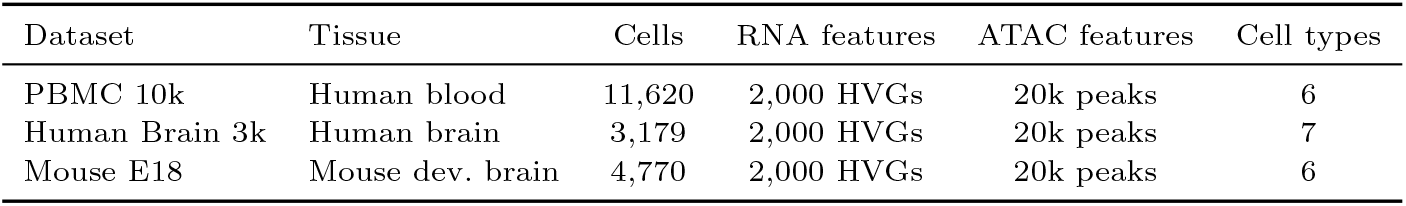
Summary of the three paired multi-omic benchmark datasets used in this study.

We compared scDisent against scVI (Lopez et al. 2018), MultiVI (Ashuach et al. 2023), scGLUE (Cao and Gao 2022), and Seurat WNN (Hao et al. 2021) (weighted nearest-neighbor). These cover RNA-only, multimodal generative, graph-guided and non-generative settings. We explicitly omit TotalVI Gayoso et al. (2021), as it is tailored for RNA and surface protein (CITE-seq) integration rather than chromatin accessibility; instead, MultiVI serves as its direct counterpart for our RNA–ATAC benchmarks.

### 2.3 Model architecture and training

scDisent contains modality-specific encoders, a dual-branch latent head, a sparse regulatory-to-expression map, and modality decoders. In the benchmark configuration, the model uses *d*_*expr*_ = 128 and *d*_*reg*_ = 128 with shared hidden dimensions (1024, 512, 256) and dropout 0.1.

The RNA encoder takes 2,000 highly variable genes, whereas the ATAC encoder takes 20,000 variable peaks and optional gene-activity features (Granja et al. 2021; Stuart et al. 2021). The modality-specific hidden states are concatenated into

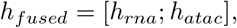

and projected to two Gaussian latent branches,

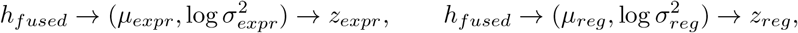

using reparameterized sampling (Kingma and Welling 2013). Branch separation is encouraged through KL regularization, a minibatch total-correlation penalty and an orthogonality term (Chen et al. 2018).

**Fig. 1.**
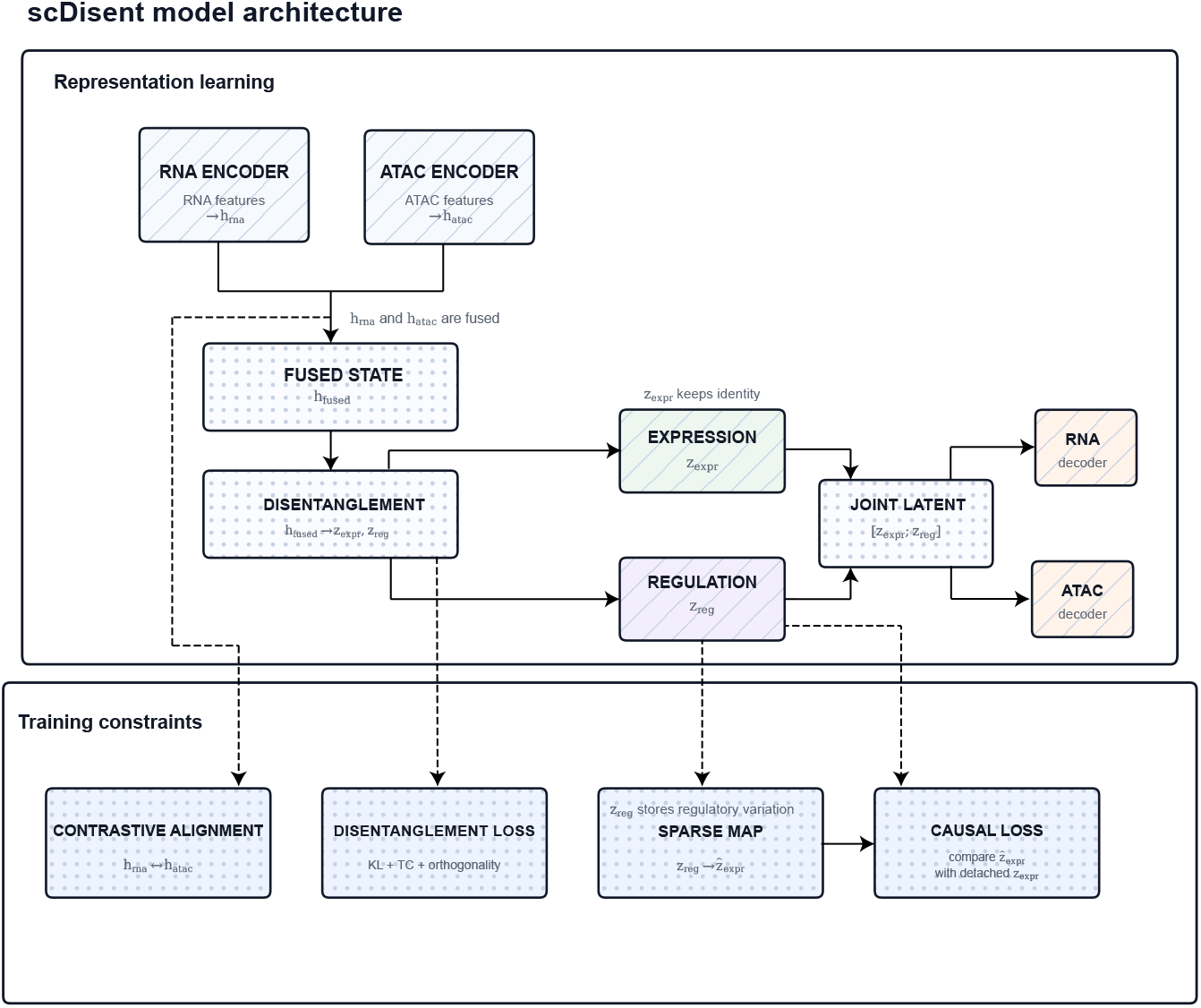
Model architecture of scDisent. Multi-omic inputs are encoded into a shared multimodal state and factorized into an expression branch and a regulatory branch. The two branches are concatenated to form the latent used by the RNA and ATAC decoders, while a smaller supervision path constrains how regulation-oriented latents predict a detached copy of the expression branch.

The model further learns a sparse mapping

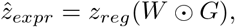

where *W* contains effect sizes and *G* is a binary gate matrix sampled with a straight-through Gumbel-Softmax estimator (Jang et al. 2016). The directional objective minimizes the distance between 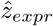 and a detached target copy of *z*_*expr*_, with additional sparsity and, when applicable, NOTEARS-style acyclicity regularization (Zheng et al. 2018). The detach operation is important because it lets the regulatory branch explain the expression branch without rewriting the identity-preserving manifold.

For reconstruction, the two branches are concatenated into

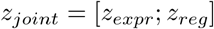

and passed to modality-specific decoders. The RNA decoder uses a zero-inflated negative binomial likelihood (Lopez et al. 2018), and the ATAC decoder uses a Bernoulli accessibility head plus an optional Gaussian gene-activity head. An auxiliary RNA decoder is applied to [*z*_*expr*_; 0] so that the expression branch remains informative on its own. Cross-modal fusion is stabilized with an InfoNCE objective between *h*_*rna*_ and *h*_*atac*_ (Oord et al. 2018).

Training uses three phases. Phase 1 learns reconstruction while the causal map is frozen. Phase 2 introduces KL, total-correlation, orthogonality, contrastive and expression-only decoder terms. Phase 3 unfreezes the causal map and optimizes the full objective. Optimization uses AdamW with cosine warm restarts, batch size 512, gradient clipping at norm 5.0, and early stopping in phase 3 (Loshchilov and Hutter 2017, 2016).

### 2.4 Evaluation

All methods were evaluated on the same processed benchmark datasets using a shared Leiden clustering sweep over thirteen resolutions from 0.1 to 2.0 (Traag et al. 2019). We report ARI and NMI as primary metrics and ASW as a compactness reference (Hubert and Arabie 1985; Vinh et al. 2010; Rousseeuw 1987). We additionally assess biological interpretability through causal-atlas inspection, lineage-specific perturbation analysis, external resource cross-checking, and quantitative branch separation.

## 3 Results

### 3.1 scDisent improves benchmark performance across immune and neural datasets

scDisent achieved the strongest ARI on all three benchmarks. The gain was largest in PBMC 10k, where scDisent reached an average ARI of 0.584 and NMI of 0.594 (all benchmark results are reported as averages across 3 random seeds). The same configuration remained competitive in Human Brain 3k (ARI 0.514, NMI 0.609) and Mouse E18 (ARI 0.299, NMI 0.372), indicating that the model does not rely on a single tissue-specific operating point.

This consistency matters because the three benchmarks differ substantially in biological geometry, spanning mature immune lineages, heterogeneous brain populations and developmental states (Chen et al. 2024; Pollen et al. 2015; Leone et al. 2015; Machold et al. 2023). The full resolution sweeps further show that the advantage is not driven by a single favorable clustering threshold.

### 3.2 The latent split separates identity-preserving and regulatory structure

Latent visualizations support the intended semantic split between the two branches. The expression branch preserves recognizable lineage geometry, whereas the regulatory branch is more diffuse and therefore less redundant with coarse population separation. In PBMC, lymphoid and myeloid populations remain sharply organized in *z*_*expr*_, while *z*_*reg*_ retains heterogeneity without reproducing the same clustering pattern (Vallejo et al. 2022; Chen et al. 2023). Supplementary baseline UMAPs show that competing methods recover reasonable manifolds, but they do not provide the same explicit separation between identity-preserving and regulation-oriented structure.

**Table 2.**
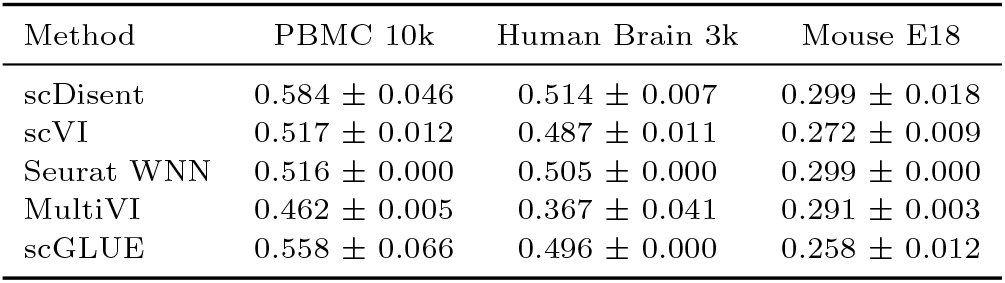
Benchmark ARI across three paired multi-omic datasets.

| Method | PBMC 10k | Human Brain 3k | Mouse E18 |
| --- | --- | --- | --- |
| scDisent | $0.584 \pm 0.046$ | $0.514 \pm 0.007$ | $0.299 \pm 0.018$ |
| scVI | $0.517 \pm 0.012$ | $0.487 \pm 0.011$ | $0.272 \pm 0.009$ |
| Seurat WNN | $0.516 \pm 0.000$ | $0.505 \pm 0.000$ | $0.299 \pm 0.000$ |
| MultiVI | $0.462 \pm 0.005$ | $0.367 \pm 0.041$ | $0.291 \pm 0.003$ |
| scGLUE | $0.558 \pm 0.066$ | $0.496 \pm 0.000$ | $0.258 \pm 0.012$ |

**Fig. 2.**
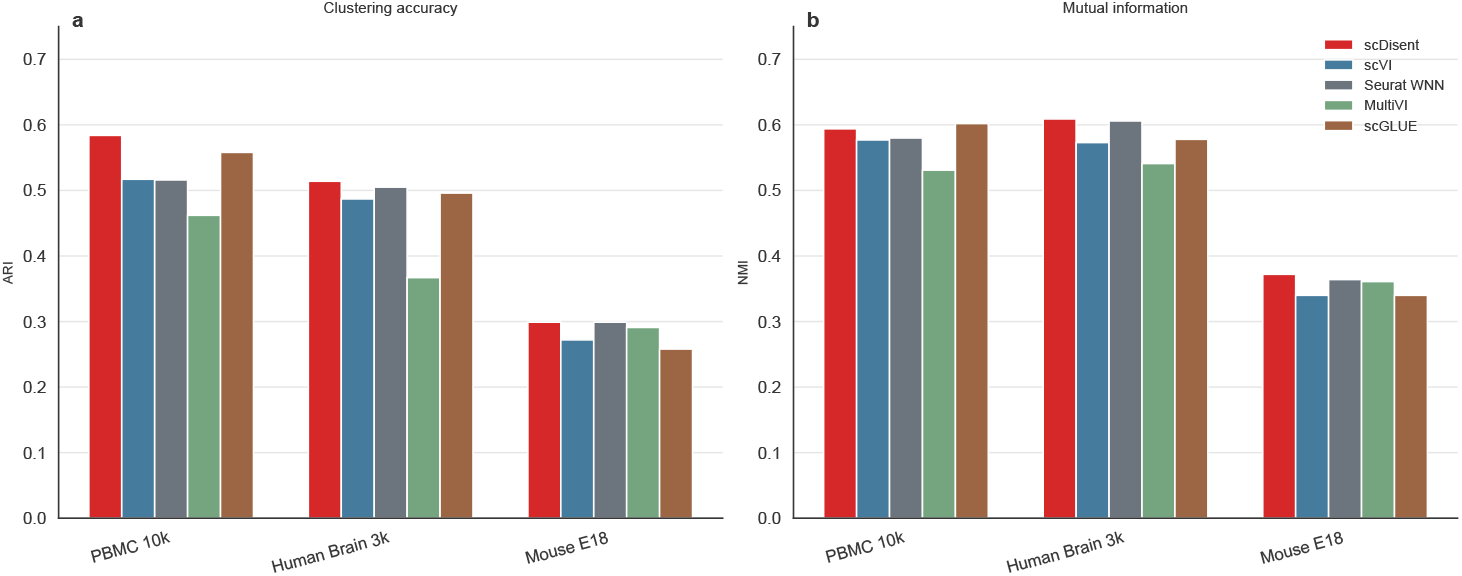
Cross-dataset benchmark performance. ARI and NMI are shown for PBMC 10k, Human Brain 3k and Mouse E18 under a shared Leiden resolution sweep. scDisent attains the strongest ARI across all three datasets while remaining competitive in NMI. Numerical values are reported in Table 2.

### 3.3 Ablations identify disentanglement as the dominant contributor

The ablation study identifies disentanglement as the main driver of performance across datasets. To ensure a strict controlled comparison, these ablation experiments were evaluated on a single seed (seed 42) for all variants, and compared against the identical single-seed full model. In PBMC 10k, the full model reaches ARI 0.627; removing disentanglement causes the largest drop, reducing ARI to 0.514, while removing causal mapping or detach-based protection reduces ARI to 0.564 and 0.565, respectively. Similar trends were observed in Mouse E18, where the full model reaches ARI 0.313 and removing disentanglement drops it to 0.291. In Human Brain 3k, the metrics were more tightly clustered across variants, but overall the results indicate that branch separation is the dominant structural contributor to scDisentś performance.

**Table 3.**
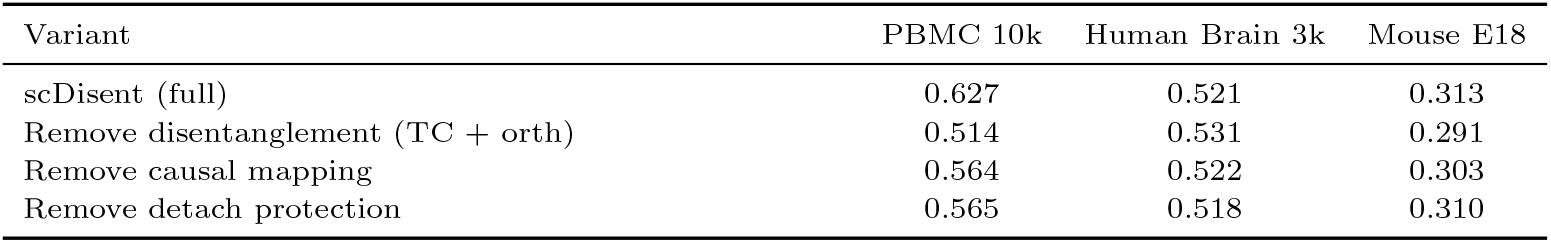
Ablation study for the core scDisent components across all datasets (single seed evaluation). Eliminating the disentanglement interface yields the largest performance drop in PBMC, confirming its structural necessity.

**Fig. 3.**
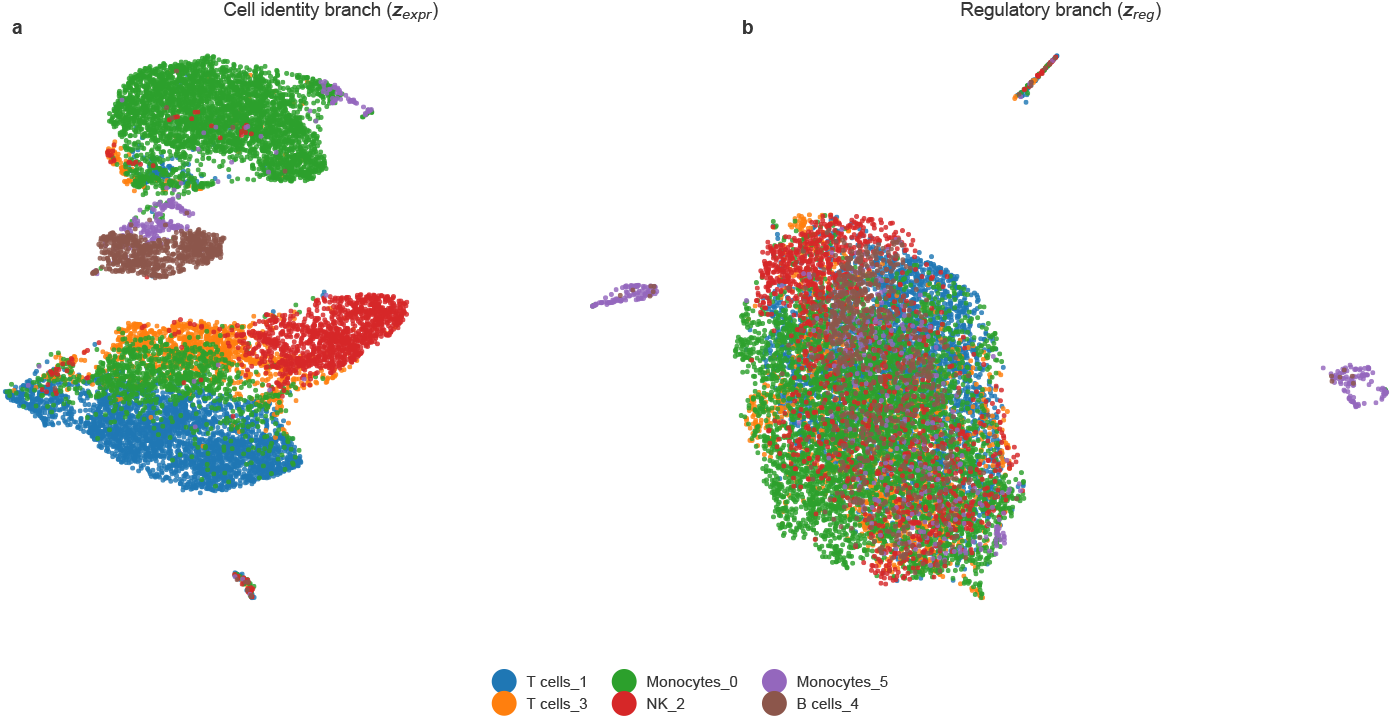
PBMC latent space visualization. The expression branch preserves clear lineage structure, whereas the regulatory branch is more diffuse and less redundant with coarse population separation. Full cross-dataset and baseline UMAP panels are provided in the supplementary material.

### 3.4 The learned regulatory atlas is sparse and structured

The learned regulatory atlas provides a compact summary of how the regulatory branch relates to expression-associated structure. In PBMC, 1,769 of 16,384 possible regulatory-to-expression links remain active, corresponding to an active-edge ratio of 10.8%. The dominant outgoing regulators are concentrated in a limited subset of latent dimensions, arguing against the interpretation that the causal layer is merely a diffuse post hoc transform.

### 3.5 Perturbation analyses recover lineage-restricted programs

Perturbation analysis on PBMC shows that the regulatory branch captures lineage-restricted immune programs rather than a generic nuisance signal. In B cells, the dominant regulator *z*_*reg* 30_ was linked to shifts in *BACH2, CD79A, MS4A1, CD74* and *HLA-DRA*, consistent with a B-cell and antigen-presentation program (Muto et al. 2010). In NK cells, *z*_*reg* 29_ was associated with canonical cytotoxic markers including *CCL5, NKG7, GZMH* and *CTSW*. In monocytes, dominant regulators were linked to *VCAN, LYZ* and *AOAH*, consistent with myeloid-state programs (Vallejo et al. 2022; Chen et al. 2023).

**Fig. 4.**
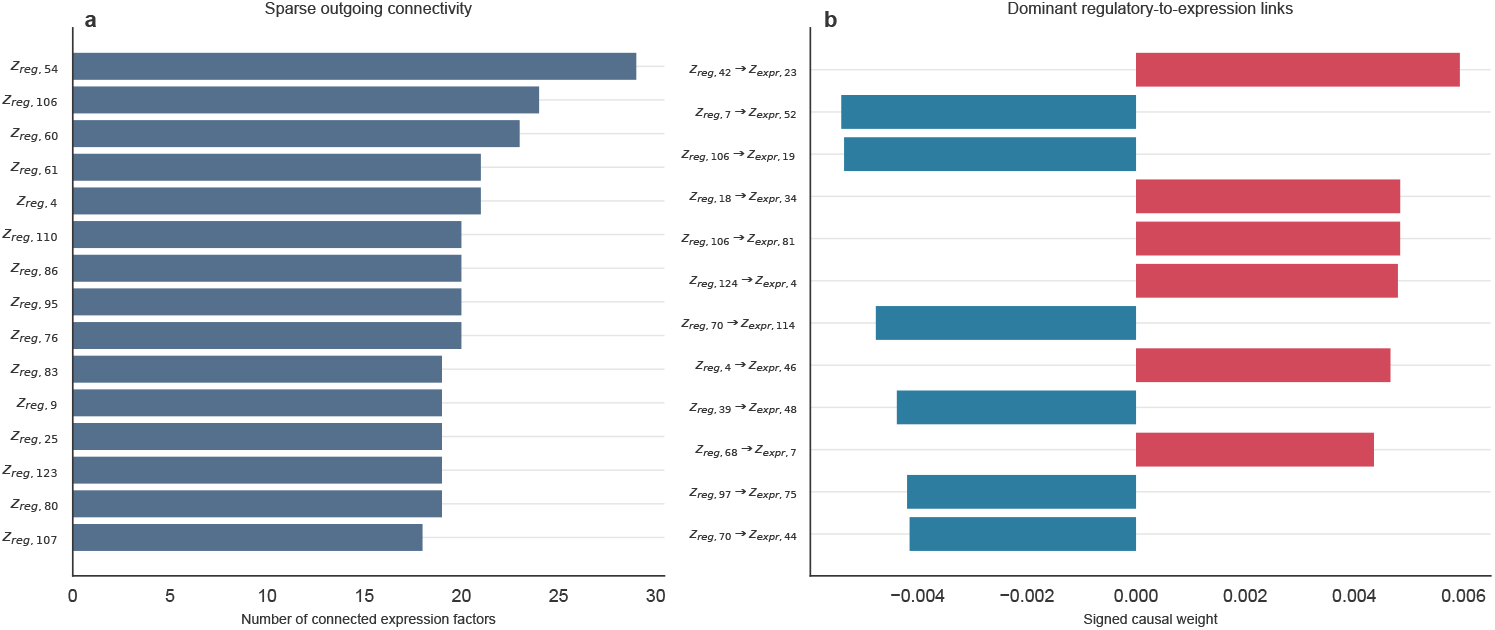
Sparse directed regulatory mapping learned in PBMC. Left, regulatory factors are ranked by the number of linked expression factors. Right, the strongest regulatory-to-expression links illustrate that only a limited subset of latent interactions dominates the map.

**Fig. 5.**
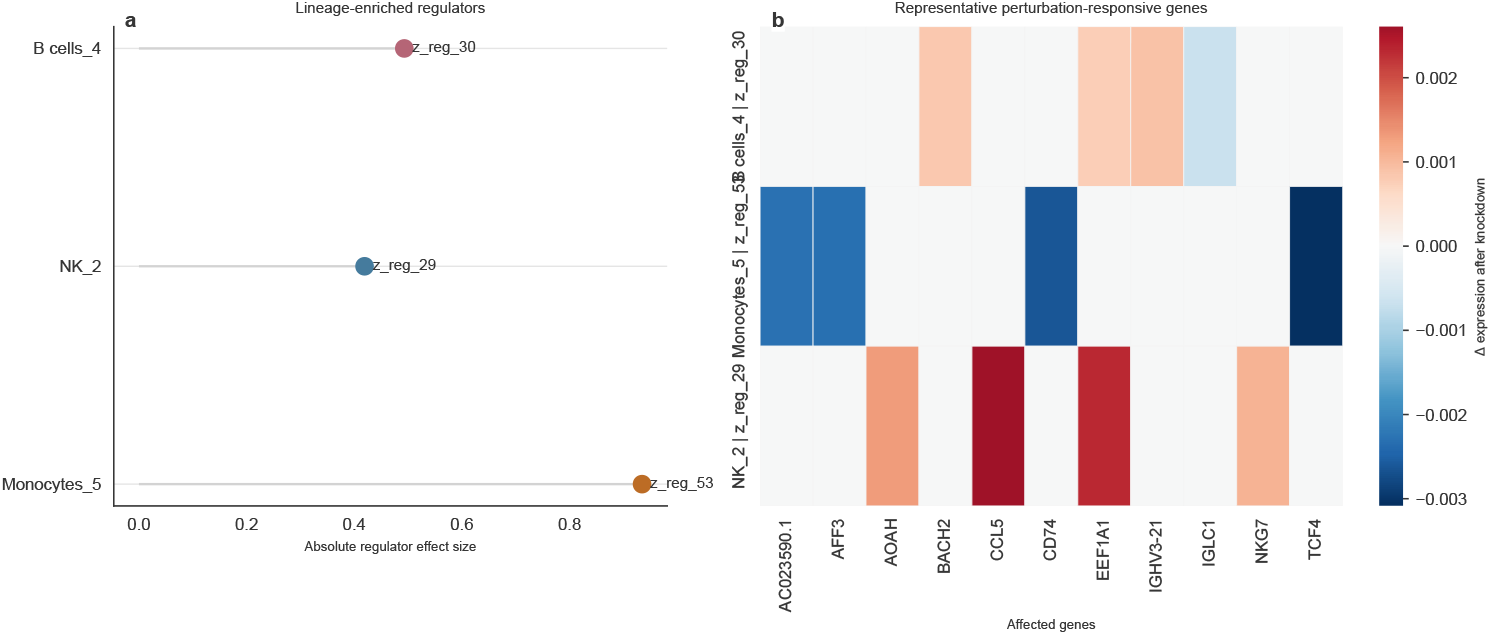
PBMC perturbation analysis. Left, lineage-enriched regulatory factors identified in B cells, NK cells and monocytes. Right, representative gene-expression shifts after in silico knockdown of those factors.

The same workflow also produced structured outputs in Human Brain and Mouse E18. Brain regulators highlighted astrocytic, excitatory-neuron and oligodendrocyte programs consistent with atlas-level distinctions (Chen et al. 2024). In Mouse E18, perturbation signatures involved *Adarb2, Erbb4, Foxp2, Satb2, Unc5d, Tnc* and *Fabp7*, matching inhibitory, projection-neuron and radial-glia-associated developmental programs (Pollen et al. 2015; Leone et al. 2015; Machold et al. 2023). These findings should be interpreted as perturbation-oriented hypotheses rather than validated mechanisms, but they show that scDisent supports regulator-centered questions across immune, neural and developmental settings.

**Fig. 6.**
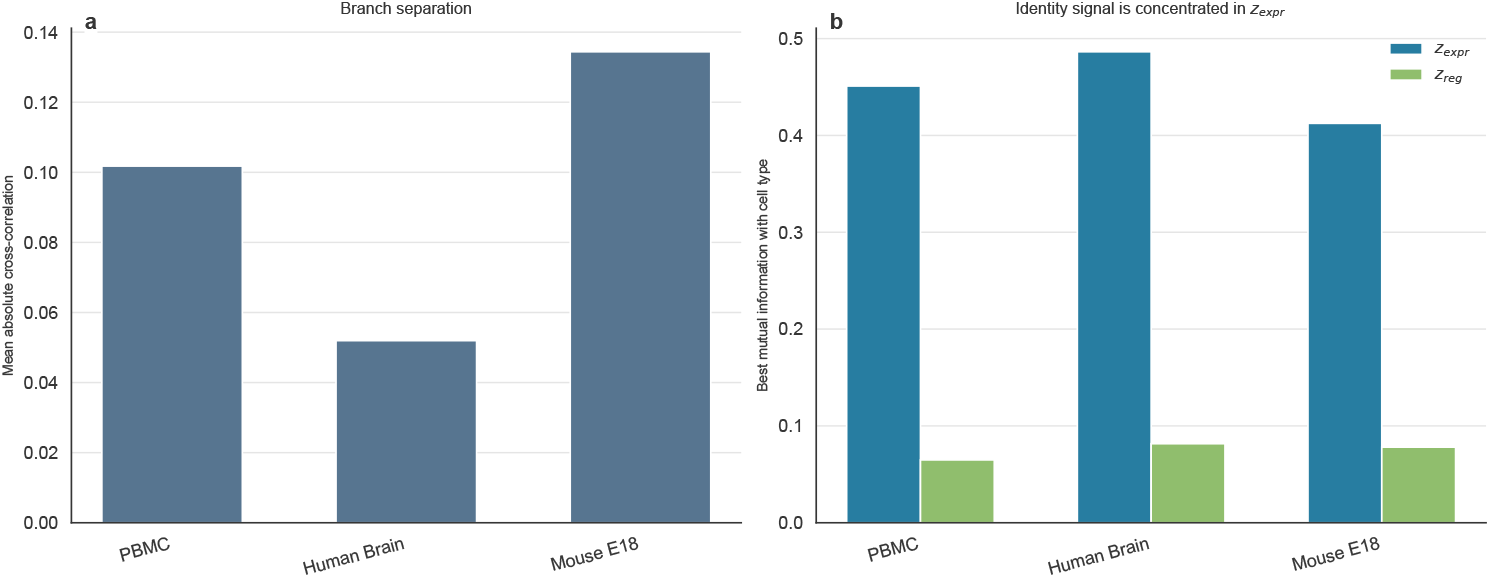
Quantitative summary of latent disentanglement across datasets. Cross-branch correlation remains modest, whereas benchmark-label information is concentrated in the expression branch rather than the regulatory branch.

### 3.6 Quantitative disentanglement supports branch separation

The disentanglement analysis provides a direct quantitative readout of branch separation. Mean absolute cross-block correlation between *z*_*expr*_ and *z*_*reg*_ remained modest across datasets: 0.102 for PBMC, 0.052 for Human Brain, and 0.134 for Mouse E18. At the same time, the strongest mutual-information score between an expression latent and the benchmark labels was much larger than the strongest score in the regulatory branch: 0.451 versus 0.065 in PBMC, 0.486 versus 0.082 in Human Brain, and 0.412 versus 0.078 in Mouse E18.

This gap indicates that identity information concentrates in *z*_*expr*_, whereas *z*_*reg*_ retains a weaker but structured residual signal. That separation matters because it argues that the perturbation results are not simply duplicated clustering information.

### 3.7 Sensitivity analysis supports the default configuration

Sensitivity scans across latent dimensionality and loss weights showed that the default configuration is close to a strong compromise across datasets. For PBMC, lower directed mapping weight (*λ*_*causal*_ = 0.1) remained competitive with ARI 0.604, whereas larger latent size (256/256) reduced ARI to 0.493. In Human Brain, nearby settings remained close to the default, with ARI values between 0.503 and 0.520. In Mouse E18, the search surface was relatively flat, with the best tested alternative reaching ARI 0.321, only slightly above the default 0.299.

## 4 Discussion

Most current multi-omic single-cell methods are optimized for integration quality, not for mechanistic decomposition (Ashuach et al. 2023; Cao and Gao 2022; Hao et al. 2021). scDisent instead changes the structure of the representation itself by separating identity-like expression variables from regulation-oriented variables and coupling them through a sparse directed interface.

The results support this view from three directions. First, scDisent remains competitive across immune, neural and developmental benchmarks, so the added structure does not sacrifice integrative performance. Second, the ablations across multiple datasets show that the main gain comes from enforcing branch semantics, with disentanglement producing the largest drop when removed. Third, the sparse atlas and perturbation analyses show that the regulatory branch remains biologically readable after training. Existing models can return strong integrated manifolds, but they do not naturally provide a branch that can be interpreted as a candidate regulatory interface (Argelaguet et al. 2020; Ashuach et al. 2023; Cao and Gao 2022; Gong et al. 2021; Korsunsky et al. 2019).

The main limitation is that the causal layer should be interpreted as a structured hypothesis space rather than proof of causal mechanism. The present study also focuses on transcriptome-accessibility datasets with matched modalities, Future work should evaluate partially observed or unpaired settings, extend validation in developmental systems, and test the same representational logic in protein-aware and time-resolved multi-omic settings (Wu et al. 2021; Zhang et al. 2022; Lee et al. 2023; Stoeckius et al. 2017; Gayoso et al. 2021; La Manno et al. 2018).

## 5 Conclusion

scDisent introduces a disentangled representation framework with directed regulatory mapping for single-cell multi-omics. By splitting the latent space into expression and regulatory branches and coupling them through a sparse directed mapping, the model improves both integration performance and biological interpretability. Across PBMC, Human Brain and Mouse E18 (10x Genomics 2020b, a, 2021), scDisent achieved the strongest benchmark clustering performance among the tested methods while producing interpretable perturbation-oriented outputs and quantitative evidence of latent separation.

## Declaration of generative AI and AI-assisted technologies in the writing process

During the preparation of this work the author used Google Gemini (AI coding assistant) in order to assist with code development, evaluation pipeline implementation, and manuscript formatting. After using this tool, the author reviewed and edited the content as needed and takes full responsibility for the content of the publication.

## Declarations

### Ethics approval and consent to participate

Not applicable. This study used only publicly available, previously released single-cell multi-omic datasets and involved no new human or animal sample collection by the author.

### Consent for publication

Not applicable.

### Availability of data and materials

The raw single-cell multi-omic datasets used in this study were originally obtained from the publicly available 10x Genomics Dataset repository. Specifically, the following filtered feature barcode matrix HDF5 files were downloaded from the respective dataset pages:

- PBMC raw dataset (10k PBMCs from a Healthy Donor): downloaded `pbmc_granulocyte_sorted_10k_filtered_feature_bc_matrix.h5` from https://www.10xgenomics.com/datasets/pbmc-from-a-healthy-donor-granulocytes-removed-through-cell-sorting-10-k-1-standard-1-0-0
- Human Brain raw dataset (3k Adult Human Brain): downloaded `human_brain_3k_filtered_feature_bc_matrix.h5` from https://www.10xgenomics.com/datasets/frozen-human-healthy-brain-tissue-3-k-1-standard-1-0-0
- Mouse E18 Brain raw dataset (5k E18 Mouse Brain): downloaded `e18_mouse_brain_fresh_5k_filtered_feature_bc_matrix.h5` from https://www.10xgenomics.com/datasets/fresh-embryonic-e-18-mouse-brain-5-k-1-standard-2-0-0

Processed paired AnnData objects derived from these raw datasets, benchmark outputs, and all code used for preprocessing, model training, and evaluation are available at https://github.com/xiguoren/scDisent.

## Authors’ contributions

G.X. conceived the study, implemented the computational framework, performed the analyses, generated the figures and tables, and wrote the main manuscript. G.X. reviewed the manuscript.

## Acknowledgements

Not applicable.

