## Supplementary Information for "scDisent: regulatory-aware disentangled representation learning for multi-omic single-cell analysis"

Guoren Xi[1*]

[1*]Datenintegrationszentrum, Universitätsklinikum Jena, Stoystr. 3, 07743, Jena, Germany.

#### 1 Supplementary methods

##### 1.1 S1. Data preprocessing

All datasets were processed into RNA and ATAC feature matrices before model fitting. In the current benchmark setting, these matrices were paired at the cell level. The RNA branch used 2,000 highly variable genes after standard quality control, while the ATAC branch used the top 20,000 peaks and additional gene-activity-style features in the processed benchmark inputs (Granja et al. 2021; Stuart et al. 2021). The unified configuration also enables motif-related ATAC features in the preprocessing pipeline, following common motif-deviation analysis strategies for single-cell chromatin accessibility data (Schep et al. 2017). Dataset-level quality-control thresholds in the default configuration included a minimum of 3 cells per feature, a minimum of 200 detected genes per cell, and a maximum mitochondrial fraction of 20%.

For RNA, normalized expression was used as encoder input, whereas raw counts were retained for the RNA reconstruction loss so that the decoder could be trained with a count-aware zero-inflated negative binomial objective (Lopez et al. 2018). For ATAC, the encoder consumed processed peak features and optional gene-activity covariates, while the decoder reconstructed binary peak accessibility logits and continuous gene-activity values when available (Granja et al. 2021; Stuart et al. 2021). This distinction between encoder inputs and decoder targets is important because it allows stable optimization without discarding the probabilistic interpretation of the reconstruction terms.

##### 1.1.1 S1.1 Dataset preparation workflow

The benchmark datasets were downloaded from 10x Genomics multiome releases and split into RNA and ATAC matrices from the joint filtered feature-barcode matrix (10x Genomics 2020b,a, 2021). RNA and ATAC modalities were then preprocessed separately and realigned by shared cell barcode before model fitting. RNA preprocessing applied cell and feature filtering, mitochondrial-content filtering, library-size normalization and highly variable gene selection. ATAC preprocessing applied feature filtering and retained the most variable peaks used in the benchmark configuration. After preprocessing, the paired modalities were restricted to the shared barcode intersection so that each training example corresponded to matched RNA and ATAC measurements from the same cell.

##### 1.1.2 S1.2 Reference label generation

The benchmark labels used for ARI and NMI were not imported as a separately curated annotation table. Instead, they were generated during preprocessing by a joint-modality labeling routine implemented in the project pipeline. Concretely, the ATAC matrix was first projected with PCA and clustered with a Leiden graph (Traag et al. 2019) to define modality-aware chromatin neighborhoods. Those chromatin-defined clusters were then mapped to biological labels by scoring RNA marker sets within the matched cells and assigning each cluster to the marker program with the highest mean expression. The resulting labels therefore combine ATAC-derived neighborhood structure with RNA-derived marker interpretation, and are saved as the `cell_type` field in the processed RNA object used for evaluation. The dataset-specific label families used in this procedure are summarized in Supplementary Table S2.

These labels act as a consistent internal reference across methods and datasets, not as a fully manual cell-atlas reannotation. In PBMC, the marker dictionary distinguishes broad immune populations such as T cells, B cells, monocytes, NK cells and dendritic cells (Vallejo et al. 2022; Chen et al. 2023). In Human Brain and Mouse E18, the marker dictionaries distinguish broad neural and developmental populations such as excitatory neurons, inhibitory neurons, astrocytes, oligodendrocytes, radial glia and related progenitor-like states (Chen et al. 2024; Pollen et al. 2015; Leone et al. 2015; Machold et al. 2023). Labels such as *Glutamatergic_0* or *Oligos_6* therefore reflect a biological label assigned to a specific Leiden-defined chromatin cluster.

##### 1.1.3 S1.3 Dataset-specific label dictionaries

The marker dictionaries used in this procedure were dataset aware. For PBMC, the labeling routine distinguished broad immune populations including T cells, B cells, monocytes, NK cells and dendritic cells using canonical marker sets such as *CD3D/IL7R*, *MS4A1/CD19*, *CD14/LYZ*, and *GNLY/NKG7* (Vallejo et al. 2022; Chen et al. 2023). For Human Brain, the dictionary separated broad neural populations including excitatory neurons, inhibitory neurons, astrocytes, oligodendrocytes, oligodendrocyte precursor cells and microglia, following broad atlas-level distinctions reported in human brain studies (Chen et al. 2024). For Mouse E18, the dictionary emphasized developmental populations such as radial glia, intermediate progenitors, glutamatergic neurons, GABAergic neurons, astrocyte-like states and oligodendrocyte-lineage states (Pollen et al. 2015; Leone et al. 2015; Machold et al. 2023).

These dictionaries were not intended to provide fine-grained atlas annotation. Their role was to generate a stable benchmark reference that is consistent with the tissue context and compatible with the joint RNA-ATAC clustering procedure. The naming convention therefore combines a broad biological label with a Leiden cluster identifier. Labels such as *T cells_1*, *Excitatory_2* or *Glutamatergic_3* should be read as automatically generated reference labels for evaluation, not as claims of exhaustive manual annotation.

###### 1.1.4 S1.4 Saved processed objects

After preprocessing, each dataset is stored as paired processed AnnData objects for RNA and ATAC together with the derived evaluation labels. The processed RNA object retains normalized encoder input, raw count layers for RNA reconstruction, and the `cell_type` field used for benchmark evaluation. The processed ATAC object retains the filtered peak representation used by the encoder together with the paired cell ordering. In addition, the pipeline stores a label array in the processed dataset directory for downstream evaluation scripts. This organization ensures that training, visualization and benchmarking all refer to the same paired cell order and the same internally generated reference labels.

###### 1.1.5 S1.5 Dataset-specific preprocessing remarks

The three benchmarks share one preprocessing framework, but their biological interpretation differs. PBMC is the most discrete setting, and its automatically generated labels mostly correspond to mature immune populations with relatively clear marker boundaries (Vallejo et al. 2022; Chen et al. 2023). Human Brain contains broader glial and neuronal classes, so the automatically generated labels should be interpreted as atlas-style major populations rather than exhaustive subtype calls (Chen et al. 2024). Mouse E18 is the most transitional setting, and its labels reflect developmental programs that are intentionally broad (Pollen et al. 2015; Leone et al. 2015; Machold et al. 2023). This difference is relevant for the downstream analyses: a label such as *Glutamatergic_2* in Mouse E18 should be read as a benchmark reference class rooted in a chromatin neighborhood and marker-supported transcriptomic identity, not as a complete developmental annotation.

##### 1.2 S2. Detailed architecture and optimization settings

All experiments used a shared backbone configuration with $d_{expr} = 128$, $d_{reg} = 128$, encoder hidden dimensions of 1024, 512 and 256, and dropout 0.1. The model configuration additionally specified causal sparsity 0.05, Gumbel temperature 1.0, and an active DAG regularization flag in the unified benchmark setting. Optimization used batch size 512, learning rate $10^{-3}$, weight decay $10^{-5}$, cosine warmup scheduling (Loshchilov and Hutter 2016) and phased training for 50, 150 and 150 epochs respectively. The default loss weights were $\beta_{KL} = 5 \times 10^{-4}$, $\gamma_{TC} = 5 \times 10^{-4}$, $\lambda_{orth} = 1.0$,

$\lambda_{causal} = 0.5$, $\lambda_{DAG} = 0.1$, $\lambda_{contrastive} = 2.5$ and $\lambda_{expr} = 1.0$. Early stopping monitored validation loss with patience 80.

Both encoders and both decoders use multilayer perceptrons with batch normalization and ReLU nonlinearities. The disentanglement head applies separate one-layer projections to the fused hidden state before computing posterior means and log-variances for the two latent branches. The causal layer contains one learnable effect matrix $W$ and one learnable gate-logit matrix of the same size. During training, the binary edge-selection process is approximated with a straight-through Gumbel-Softmax sampler (Jang et al. 2016); during evaluation, adjacency is read out by thresholding the gate probabilities at 0.5. The auxiliary $z_{expr}$-only decoder is architecturally identical to the main RNA decoder and differs only in receiving $[z_{expr}; 0]$ instead of the full latent vector.

Gradient handling is part of the method rather than an implementation detail. In the default model, the causal branch predicts a detached copy of $z_{expr}$, which prevents the prediction target from being dragged by the causal objective. The ablation without detach removes exactly this barrier and therefore tests whether the directed map can be optimized without corrupting the identity-preserving manifold.

The implementation also uses phase-local warmup coefficients. During phase 2, KL, TC, orthogonality, contrastive alignment and the auxiliary decoder are gradually ramped over the first 30 local phase epochs. During phase 3, the causal loss is ramped over the first 20 local phase epochs. These schedules match the loss implementation used in training and are part of the default configuration rather than post hoc tuning.

##### 1.2.1 S2.1 Loss scheduling and annealing

KL regularization is annealed globally over the first 50 epochs according to $\alpha_{KL}(e) = \min(1, e/50)$, while the phase-local warmup controls how strongly the branch-separation terms are introduced after reconstruction pretraining. In phase 2, $\alpha_{phase2}(e_2) = \min(1, e_2/30)$ gates TC, orthogonality, contrastive alignment and the auxiliary decoder. In phase 3, $\alpha_{causal}(e_3) = \min(1, e_3/20)$ gates the causal and sparsity-related terms. This combination is important because KL collapse and unstable total-correlation estimates are both more likely when regularization is applied too aggressively before the encoder-decoder backbone has stabilized (Kingma and Welling 2013; Chen et al. 2018). The supplementary ablation tables should therefore be read in the context of this schedule rather than as isolated weight changes.

##### 1.2.2 S2.2 Freeze and unfreeze policy

The trainer freezes the causal-map parameters during phases 1 and 2 and only unfreezes them in phase 3. All other parameters remain trainable throughout the later stages so that the encoders, disentanglement head and decoders can co-adapt when the latent semantics become more constrained. This choice differs from a more naive pretrain-then-freeze design and was used to avoid posterior collapse or brittle latent separation when KL and TC terms were introduced.

#### 1.3 S3. Evaluation details

For every model and dataset, Leiden clustering was evaluated over a fixed resolution grid spanning 0.1, 0.15, 0.2, 0.25, 0.3, 0.4, 0.5, 0.6, 0.8, 1.0, 1.2, 1.5 and 2.0. Reported scores correspond to the best value within this grid. This protocol was applied consistently to scDisent and all baselines in order to minimize resolution-selection bias. The best resolutions differed substantially across methods and datasets, which confirms that reporting a single fixed clustering resolution would have been misleading.

The evaluation suite also included latent-space visualization, branch-separation summaries, causal adjacency extraction, lineage-enriched regulator ranking and in silico perturbation summaries. These analyses were performed on trained checkpoints rather than during model selection. Clustering metrics were used to compare integration quality, whereas the causal-atlas and perturbation analyses were used to assess whether the learned representation remained biologically interpretable after training.

##### 1.3.1 S3.1 Visual benchmark interpretation

UMAP visualizations (McInnes et al. 2018) were used as qualitative companions to the quantitative benchmark rather than as standalone evidence. For scDisent, three views were inspected for each dataset: $z_{expr}$, $z_{reg}$ and the combined latent state. For the reference methods, the single latent manifold was visualized directly. This evaluation design matters because the biological claim of the paper is not that scDisent produces universally prettier UMAPs, but that it returns a pair of complementary latent views with distinct intended semantics. The supplementary baseline UMAP figures should therefore be read as a control showing that standard baselines can yield reasonable manifolds without yielding an explicit separation between identity-preserving and regulation-oriented structure.

#### 1.4 S4. Ablation settings

The ablation study evaluated three targeted removals of the full architecture across all three datasets (PBMC 10k, Human Brain 3k, Mouse E18): removal of disentanglement regularization by setting $\gamma_{TC} = 0$ and $\lambda_{orth} = 0$; removal of causal mapping by setting $\lambda_{causal} = 0$; and removal of gradient isolation by disabling the detach-based protection in the causal loss. These ablations were chosen to match the three main architectural claims in the paper: that branch separation is necessary, that the directional regulatory interface contributes beyond ordinary reconstruction, and that the causal interface must be prevented from overwriting the identity-preserving branch. To ensure a strict controlled comparison, these ablation experiments were evaluated on a single seed (seed 42) for all variants, whereas the main benchmark results report robust averages across three random seeds.

##### 1.4.1 S4.1 Full ablation interpretation

The ablation suite is intentionally compact because each setting is meant to isolate one conceptual failure mode rather than to exhaustively enumerate all combinations of removed losses. Removing disentanglement collapses the core branch semantics.

Removing the causal map tests whether explicit regulatory-to-expression structure contributes beyond a two-branch VAE. Removing the detach barrier tests whether the same directional objective remains useful once causal gradients are allowed to corrupt the target branch. These settings therefore function as architectural controls rather than as generic hyperparameter perturbations.

#### 1.5 S5. Interpretation pipeline

Regulatory interpretation was carried out by extracting the learned latent causal adjacency matrix and lineage-enriched regulatory dimensions from trained scDisent models for PBMC, Human Brain and Mouse E18. Perturbation summaries were generated by in silico knockdown of candidate regulatory dimensions followed by inspection of the strongest gene-expression shifts within each lineage or cell state. The resulting outputs were used to identify candidate immune, neural and developmental programs associated with the major annotated populations in each dataset. For PBMC, we additionally cross-checked perturbation signatures against the DoRothEA resource (Garcia-Alonso et al. 2019) using the evaluation script `scripts/eval/run_dorothea_validation.py`; this analysis is included as a supporting resource-based validation rather than as proof of mechanism.

More concretely, candidate regulators were prioritized by lineage enrichment within the regulatory branch, and perturbation effects were summarized by ranking the largest predicted gene-expression changes after targeted suppression of the selected regulatory dimension. These outputs were then compared with known lineage markers and curated resources when available. We emphasize that this procedure is intended to surface candidate regulatory programs rather than to claim direct identification of causal molecular mechanisms.

##### 1.5.1 S5.1 Regulator prioritization

Within each dataset, regulator candidates were ranked by their lineage enrichment and by the magnitude of their downstream perturbation signatures. This strategy does not assume that every regulatory latent corresponds to a single transcription factor or pathway. Instead, it treats each latent dimension as a compact regulatory program whose interpretability must be assessed empirically through enrichment and perturbation summaries.

##### 1.5.2 S5.2 Biological cross-checking

When curated prior knowledge was available, perturbation outputs were compared with known lineage markers and public resources such as DoRothEA (Garcia-Alonso et al. 2019). These comparisons were used as plausibility checks rather than as strict labels for the latent variables. This distinction is important because the biological interpretation of latent factors is many-to-many and because the current study is based on observational multi-omic measurements rather than direct perturbation readouts.

##### 1.6 S6. Sensitivity analysis design

Sensitivity analysis scanned latent dimensionality (64/64, 128/128, 256/256), causal weight ($\lambda_{causal}$ = 0.1, 0.5, 1.0, 1.5), and contrastive weight ($\lambda_{contrastive}$ = 0.1, 0.5, 1.0, 2.5) across all three datasets. These scans were intended to test whether the unified benchmark configuration is unusually fragile and to provide a local stability map around the default setting.

###### 1.6.1 S6.1 Why these hyperparameters were scanned

These three hyperparameter groups correspond to the main non-trivial design choices of scDisent. Latent dimensionality determines how much capacity is assigned to the identity-preserving and regulation-oriented branches. The causal weight determines how strongly the sparse directional interface is enforced relative to reconstruction and disentanglement. The contrastive weight determines how strongly modality-specific hidden states are aligned before branch factorization. Together, these scans cover the most plausible sources of overfitting or under-constraint in the default configuration.

##### 1.7 S7. Computational considerations

The current configuration prioritizes representational quality and interpretability over minimal compute. The main added costs relative to a conventional single-latent VAE come from total-correlation estimation, the second latent branch, and the causal regularization terms. Although the current study does not present a full wall-clock benchmark against all baselines, these components make scDisent a richer but heavier model than purely integration-oriented alternatives.

Among these components, total-correlation estimation is the most computationally distinctive because it requires minibatch-wise density comparisons across latent samples, while the causal layer introduces additional parameters and warmup logic beyond a standard multimodal VAE. The method therefore targets settings in which interpretability and biological utility justify a modest increase in training complexity.

###### 1.7.1 S7.1 Complexity summary

Let $B$ denote batch size, $d = d_{expr} + d_{reg}$ the total latent dimension, $d_h$ the encoder hidden width, $p$ the RNA feature dimension and $q$ the ATAC feature dimension. The dominant feed-forward cost per batch is linear in the feature and hidden sizes of the encoder-decoder backbone and is comparable to that of a standard multimodal VAE with the same widths. Relative to a single-latent baseline, the extra costs introduced by scDisent arise from three sources:

$$\mathcal{O}(Bd_h d) \quad \text{for the second latent branch,}$$

$$\mathcal{O}(B^2 d) \quad \text{for minibatch-based total-correlation estimation,}$$
$$\mathcal{O}(d_{expr} d_{reg}) \quad \text{for the sparse causal map and its penalties.}$$

The total-correlation term is therefore the only component with explicit quadratic dependence on batch size in the current implementation, whereas the causal map scales with latent dimensionality rather than with the number of observed genes or peaks.

##### 1.7.2 S7.2 Practical compute interpretation

In practical terms, scDisent should be understood as modestly heavier than a conventional embedding-oriented multimodal model, although it remains far lighter than methods that attempt graph-level inference over observed features. This computational tradeoff is deliberate: the method spends extra budget to obtain a representation with more explicit branch semantics and a sparse regulator-to-expression interface. For the benchmark sizes studied here, that added cost remained acceptable for repeated ablation, sensitivity and discovery analyses.

### 2 Supplementary results overview

The supplementary material documents the quantitative and interpretive record underlying the main-text claims. It serves four purposes. First, it reports detailed ablation and sensitivity settings that are summarized only briefly in the main Results section. Second, it provides complete Leiden resolution sweeps so that benchmark comparisons can be inspected beyond the best-performing clustering resolution. Third, it extends the visualization record with dataset-specific UMAPs, baseline latent UMAPs and full disentanglement panels. Fourth, it records supporting biological interpretation outputs, including cross-dataset discovery summaries and the DoRothEA-based PBMC cross-check (Garcia-Alonso et al. 2019).

These supplementary analyses matter because several of the paper's claims depend on structure across many related results rather than on one isolated figure. For example, the claim that scDisent improves clustering robustness is supported not only by the peak ARI table but also by the full resolution sweeps across methods and datasets. Likewise, the claim that disentanglement improves biological interpretability is supported not only by the main PBMC figures but also by the supplementary branch-separation analyses, cross-dataset discovery outputs and baseline UMAP comparisons (Argelaguet et al. 2021; Zhu et al. 2023).

The supplement should therefore be read as the supporting record for the study rather than as miscellaneous overflow.

#### 2.1 Supplementary benchmark robustness

Supplementary Tables S8, S9 and S10 report the full Leiden resolution sweeps for all methods and datasets. These results show that the benchmark story in the main text is not driven by one favorable clustering threshold. Instead, they document how each method behaves over a shared evaluation grid and make clear that the relative ranking varies less by lucky resolution choice than by the quality of the underlying latent geometry.

Across datasets, these sweeps also show that different reference methods prefer substantially different operating points. This is precisely why the unified sweep protocol is needed: fixing one clustering resolution across all methods would hide genuine differences in manifold organization and would overstate or understate performance depending on the dataset. The full sweep record therefore serves as a methodological control for the benchmark, not merely as extra data.

#### 2.2 Supplementary ablation and sensitivity record

Supplementary Table S3 and Supplementary Table S7 extend the main-text architectural and hyperparameter analyses. The ablation table records the exact configuration changes used to isolate each architectural component, while the sensitivity table documents how nearby latent and loss-weight settings affect performance across tissues. Together, these tables support the claim that the full model is not only strong at one hand-tuned operating point but also interpretable as a coherent design rather than a fragile parameter combination.

The supplementary ablation record is especially useful for separating architectural interpretation from benchmark outcome. In the main text, the ablation result is summarized as evidence that disentanglement is the dominant contributor. In the supplement, the explicit configuration changes make that claim auditable: one can see which objective terms were removed, how much the score changed, and which conclusion is being drawn from each manipulation. This level of detail matters because the paper is making a design claim, not only a leaderboard claim.

#### 2.3 Supplementary visualization and interpretation record

The supplementary figures provide the full visualization record behind the main-text summaries. This includes dataset-specific disentangled UMAPs, baseline UMAP comparisons, dataset-specific branch-separation panels, and cross-dataset discovery figures. These assets complement the story-driven main-text figures and document the broader analysis footprint that supports the paper's claims about latent structure and biological interpretability (McInnes et al. 2018; Argelaguet et al. 2021; Zhu et al. 2023).

##### How to read the supplementary labels

The automatically generated benchmark labels used throughout the supplementary figures should be interpreted as reference classes for evaluation and visualization. They are most reliable as broad lineage or state descriptors, especially in PBMC and in the major neural classes of Human Brain. In Mouse E18, where transitions are common, they should be read more cautiously as marker-supported developmental neighborhoods rather than exhaustive biological annotations.

#### 2.4 Supplementary biological cross-check record

The supplementary tables and notes also document how biological interpretation claims were kept conservative. In particular, the DoRothEA summary is presented as a plausibility-oriented resource cross-check rather than as definitive regulator annotation (Garcia-Alonso et al. 2019). Likewise, the cross-dataset discovery notes are framed as evidence that regulator-centered summaries remain coherent outside PBMC. This distinction matters because the paper argues for improved biological interpretability, not for direct causal proof from observational data alone (Argelaguet et al. 2021; Zhu et al. 2023).

### 3 Supplementary tables

The supplementary tables provide the full quantitative record behind the main-text summaries, including detailed ablation settings, complete resolution sweeps, and supporting resource-based validation.

**Table S1** Summary of the three paired multi-omic benchmark datasets used in this study.

| Dataset | Tissue | Cells | RNA features | ATAC features | Cell types |
|---|---|---|---|---|---|
| PBMC 10k | Human blood | 11,620 | 2,000 HVGs | 20k peaks | 6 |
| Human Brain 3k | Human brain | 3,179 | 2,000 HVGs | 20k peaks | 7 |
| Mouse E18 | Mouse dev. brain | 4,770 | 2,000 HVGs | 20k peaks | 6 |

**Table S2** Supplementary Table. Dataset-specific reference label generation scheme used for benchmark evaluation. Broad biological classes were assigned to ATAC-defined Leiden clusters by RNA marker scoring within matched cells. Cluster suffixes denote chromatin-cluster identifiers rather than manually curated subtypes.

| Dataset | Broad label families | Marker basis |
|---|---|---|
| PBMC 10k | T cells, B cells, Monocytes, NK cells, Dendritic cells | Canonical immune markers including *CD3D/IL7R*, *MS4A1/CD19*, *CD14/LYZ*, and *GNLY/NKG7*. |
| Human Brain 3k | Excitatory neurons, Inhibitory neurons, Astrocytes, Oligodendrocytes, OPCs, Microglia | Neural and glial markers including *SLC17A7*, *GAD1/GAD2*, *GFAP/AQP4*, *PLP1/MBP*, and *PDGFRA/CSPG4*. |
| Mouse E18 | Radial glia, Intermediate progenitors, Glutamatergic neurons, GABAergic neurons, Astrocyte-like states, Oligodendrocyte-lineage states | Developmental markers including *Pax6/Slc1a3*, *Eomes*, *Slc17a7/Tbr1*, *Gad1/Gad2*, and *Olig1/Olig2/Pdgfra*. |

**Table S3** Detailed PBMC ablation configurations and quantitative outcomes.

| Variant | Configuration change | PBMC ARI (seed 42) | Δ |
|---|---|---|---|
| scDisent (full) | None | 0.627 | +0.000 |
| Remove disentanglement | gamma_TC = 0; lambda_orth = 0 | 0.514 | -0.114 |
| Remove causal mapping | lambda_causal = 0 | 0.564 | -0.063 |
| Remove detach protection | detach(Z_expr) -> Z_expr in L_causal | 0.565 | -0.062 |

**Table S4** PBMC DoRothEA cross-check for perturbation-derived regulators. This table is intended as supporting resource-based validation rather than definitive regulator identification.

| Regulator | Matched TF | Empirical p-value | Overlap |
|---|---|---|---|
| z_reg_2 | FOXA1 | 0.0050 | 10 |
| z_reg_29 | FOXA1 | 0.0050 | 10 |
| z_reg_30 | FOXA1 | 0.0050 | 10 |
| z_reg_51 | FOXA1 | 0.0050 | 10 |
| z_reg_53 | FOXA1 | 0.0050 | 10 |
| z_reg_96 | FOXA1 | 0.0050 | 10 |

**Table S5** Quantitative separation between expression and regulatory latent branches.

| Dataset | Mean \|corr\| | Max \|corr\| | Best MI ($z_{expr}$) | Best MI ($z_{reg}$) |
|---|---|---|---|---|
| PBMC | 0.102 | 0.513 | 0.451 | 0.065 |
| Brain | 0.052 | 0.364 | 0.486 | 0.082 |
| E18 | 0.134 | 0.688 | 0.412 | 0.078 |

**Table S6** Cross-dataset summary of lineage- or cell-state-enriched regulators and representative perturbation-responsive genes.

| Dataset | Cell type | Top regulator | Effect size | Representative genes |
|---|---|---|---|---|
| PBMC | Monocytes_5 | z_reg_53 | 0.935 | TCF4, CD74, AFF3 |
| PBMC | B cells_4 | z_reg_30 | -0.493 | IGHV3-21, BACH2, EEF1A1 |
| PBMC | NK_2 | z_reg_29 | -0.419 | CCL5, EEF1A1, AOAH |
| PBMC | T cells_3 | z_reg_51 | 0.317 | EEF1A1, LTB, BCL2 |
| Human Brain | Oligos_6 | z_reg_115 | 1.054 | HSPH1, LRMDA, P4HA1 |
| Human Brain | Oligos_4 | z_reg_11 | 0.424 | KCNIP4, PCDH15, CSMD1 |
| Human Brain | Excitatory_2 | z_reg_63 | -0.402 | KCNIP4, ROBO2, DPP10 |
| Human Brain | Oligos_5 | z_reg_87 | -0.392 | CEMIP2, SYTL3, DSCAM |
| Mouse E18 | GABAergic_5 | z_reg_59 | 0.544 | Adarb2, Erbb4, Gm42418 |
| Mouse E18 | Glutamatergic_3 | z_reg_102 | 0.453 | Ptprk, Dcc, Gm42418 |
| Mouse E18 | Glutamatergic_2 | z_reg_92 | -0.436 | Satb2, Unc5d, Ptn |
| Mouse E18 | Radial Glia_4 | z_reg_92 | 0.392 | Dcc, Tnc, Ptprz1 |

**Table S7** Sensitivity analysis over latent dimensionality and loss weights.

| Dataset | Setting | ARI | NMI | ASW | Resolution |
|---|---|---|---|---|---|
| PBMC 10k | causal_01 | 0.604 | 0.605 | 0.129 | 0.20 |
| PBMC 10k | causal_10 | 0.515 | 0.573 | 0.113 | 0.15 |
| PBMC 10k | causal_15 | 0.582 | 0.585 | 0.107 | 0.15 |
| PBMC 10k | contra_01 | 0.512 | 0.580 | 0.125 | 0.10 |
| PBMC 10k | contra_05 | 0.502 | 0.552 | 0.119 | 0.15 |
| PBMC 10k | contra_10 | 0.483 | 0.544 | 0.118 | 0.10 |
| PBMC 10k | dim_256 | 0.493 | 0.552 | 0.138 | 0.10 |
| PBMC 10k | dim_64 | 0.584 | 0.588 | 0.113 | 0.20 |
| Human Brain 3k | causal_01 | 0.505 | 0.611 | 0.019 | 0.10 |
| Human Brain 3k | causal_10 | 0.520 | 0.623 | 0.015 | 0.15 |
| Human Brain 3k | causal_15 | 0.507 | 0.612 | 0.013 | 0.10 |
| Human Brain 3k | contra_01 | 0.506 | 0.601 | 0.040 | 0.25 |
| Human Brain 3k | contra_05 | 0.503 | 0.596 | 0.014 | 0.25 |
| Human Brain 3k | contra_10 | 0.511 | 0.614 | 0.001 | 0.10 |
| Human Brain 3k | dim_256 | 0.500 | 0.586 | -0.001 | 0.10 |
| Human Brain 3k | dim_64 | 0.518 | 0.602 | 0.011 | 0.25 |
| Mouse E18 | causal_01 | 0.313 | 0.379 | 0.014 | 0.30 |
| Mouse E18 | causal_10 | 0.278 | 0.366 | 0.018 | 0.30 |
| Mouse E18 | causal_15 | 0.300 | 0.377 | 0.017 | 0.40 |
| Mouse E18 | contra_01 | 0.283 | 0.359 | 0.012 | 0.30 |
| Mouse E18 | contra_05 | 0.316 | 0.387 | -0.008 | 0.40 |
| Mouse E18 | contra_10 | 0.287 | 0.379 | 0.012 | 0.20 |
| Mouse E18 | dim_256 | 0.321 | 0.395 | 0.007 | 0.40 |
| Mouse E18 | dim_64 | 0.288 | 0.357 | 0.015 | 0.40 |

**Table S8** PBMC 10k Leiden resolution sweep. Best-performing operating points for each method are shown in bold.

| | scDisent | | scVI | | MultiVI | | scGLUE | | WNN | |
|---|---|---|---|---|---|---|---|---|---|---|
| Resolution | ARI | NMI | ARI | NMI | ARI | NMI | ARI | NMI | ARI | NMI |
| 0.10 | 0.357 | 0.466 | 0.359 | 0.472 | **0.476** | **0.538** | **0.516** | **0.587** | 0.492 | 0.559 |
| 0.15 | 0.511 | 0.567 | 0.500 | 0.568 | 0.400 | 0.513 | 0.504 | 0.585 | 0.495 | 0.561 |
| 0.20 | **0.627** | **0.610** | 0.523 | 0.584 | 0.401 | 0.523 | 0.477 | 0.557 | **0.516** | **0.580** |
| 0.25 | 0.480 | 0.535 | 0.465 | 0.541 | 0.403 | 0.524 | 0.380 | 0.525 | 0.458 | 0.529 |
| 0.30 | 0.577 | 0.583 | **0.583** | **0.579** | 0.409 | 0.546 | 0.386 | 0.534 | 0.411 | 0.524 |
| 0.40 | 0.436 | 0.533 | 0.544 | 0.583 | 0.372 | 0.533 | 0.407 | 0.541 | 0.404 | 0.513 |
| 0.50 | 0.370 | 0.523 | 0.543 | 0.584 | 0.304 | 0.518 | 0.396 | 0.540 | 0.448 | 0.536 |
| 0.60 | 0.319 | 0.509 | 0.540 | 0.575 | 0.230 | 0.485 | 0.388 | 0.539 | 0.455 | 0.548 |
| 0.80 | 0.250 | 0.473 | 0.447 | 0.548 | 0.184 | 0.464 | 0.321 | 0.522 | 0.413 | 0.530 |
| 1.00 | 0.236 | 0.467 | 0.429 | 0.536 | 0.159 | 0.450 | 0.301 | 0.516 | 0.321 | 0.508 |
| 1.20 | 0.172 | 0.446 | 0.332 | 0.506 | 0.147 | 0.447 | 0.270 | 0.504 | 0.308 | 0.492 |
| 1.50 | 0.131 | 0.430 | 0.272 | 0.483 | 0.129 | 0.437 | 0.215 | 0.478 | 0.298 | 0.488 |
| 2.00 | 0.120 | 0.423 | 0.236 | 0.468 | 0.109 | 0.429 | 0.175 | 0.471 | 0.247 | 0.468 |

**Table S9** Human Brain 3k Leiden resolution sweep. Best-performing operating points for each method are shown in bold.

| | scDisent | | scVI | | MultiVI | | scGLUE | | WNN | |
|---|---|---|---|---|---|---|---|---|---|---|
| Resolution | ARI | NMI | ARI | NMI | ARI | NMI | ARI | NMI | ARI | NMI |
| 0.10 | 0.507 | 0.614 | **0.514** | **0.593** | **0.392** | **0.541** | 0.484 | 0.579 | 0.452 | 0.566 |
| 0.15 | **0.521** | **0.626** | 0.493 | 0.574 | 0.280 | 0.487 | **0.500** | **0.588** | 0.475 | 0.578 |
| 0.20 | 0.493 | 0.605 | 0.481 | 0.561 | 0.277 | 0.486 | 0.500 | 0.588 | 0.494 | 0.595 |
| 0.25 | 0.494 | 0.607 | 0.488 | 0.570 | 0.293 | 0.511 | 0.494 | 0.578 | 0.494 | 0.595 |
| 0.30 | 0.494 | 0.608 | 0.501 | 0.585 | 0.267 | 0.497 | 0.354 | 0.508 | 0.494 | 0.597 |
| 0.40 | 0.403 | 0.549 | 0.501 | 0.585 | 0.250 | 0.484 | 0.369 | 0.511 | **0.505** | **0.606** |
| 0.50 | 0.359 | 0.519 | 0.495 | 0.578 | 0.218 | 0.479 | 0.324 | 0.490 | 0.501 | 0.598 |
| 0.60 | 0.359 | 0.519 | 0.383 | 0.519 | 0.209 | 0.473 | 0.312 | 0.487 | 0.501 | 0.597 |
| 0.80 | 0.266 | 0.474 | 0.348 | 0.499 | 0.183 | 0.452 | 0.264 | 0.473 | 0.492 | 0.579 |
| 1.00 | 0.265 | 0.472 | 0.248 | 0.453 | 0.187 | 0.453 | 0.265 | 0.472 | 0.333 | 0.496 |
| 1.20 | 0.238 | 0.463 | 0.220 | 0.437 | 0.176 | 0.446 | 0.223 | 0.446 | 0.308 | 0.489 |
| 1.50 | 0.190 | 0.439 | 0.195 | 0.420 | 0.143 | 0.435 | 0.209 | 0.439 | 0.207 | 0.446 |
| 2.00 | 0.169 | 0.423 | 0.179 | 0.413 | 0.124 | 0.423 | 0.168 | 0.421 | 0.174 | 0.420 |

**Table S10** Mouse E18 Leiden resolution sweep. Best-performing operating points for each method are shown in bold.

| | scDisent | | scVI | | MultiVI | | scGLUE | | WNN | |
|---|---|---|---|---|---|---|---|---|---|---|
| Resolution | ARI | NMI | ARI | NMI | ARI | NMI | ARI | NMI | ARI | NMI |
| 0.10 | 0.116 | 0.245 | 0.180 | 0.275 | 0.190 | 0.302 | 0.165 | 0.278 | 0.136 | 0.237 |
| 0.15 | 0.183 | 0.316 | 0.184 | 0.295 | 0.223 | 0.322 | 0.169 | 0.283 | 0.199 | 0.296 |
| 0.20 | 0.263 | 0.361 | 0.260 | 0.325 | **0.293** | **0.372** | 0.236 | 0.311 | 0.203 | 0.307 |
| 0.25 | 0.279 | 0.367 | 0.263 | 0.333 | 0.284 | 0.354 | 0.246 | 0.344 | 0.201 | 0.311 |
| 0.30 | **0.313** | **0.387** | **0.279** | **0.345** | 0.282 | 0.353 | **0.248** | **0.344** | **0.299** | **0.364** |
| 0.40 | 0.287 | 0.365 | 0.252 | 0.342 | 0.230 | 0.334 | 0.241 | 0.339 | 0.270 | 0.346 |
| 0.50 | 0.283 | 0.356 | 0.259 | 0.351 | 0.201 | 0.334 | 0.232 | 0.339 | 0.298 | 0.363 |
| 0.60 | 0.275 | 0.365 | 0.248 | 0.346 | 0.174 | 0.333 | 0.216 | 0.330 | 0.232 | 0.333 |
| 0.80 | 0.231 | 0.345 | 0.222 | 0.344 | 0.155 | 0.335 | 0.199 | 0.330 | 0.242 | 0.348 |
| 1.00 | 0.212 | 0.355 | 0.187 | 0.334 | 0.150 | 0.342 | 0.194 | 0.343 | 0.198 | 0.342 |
| 1.20 | 0.216 | 0.355 | 0.165 | 0.325 | 0.126 | 0.340 | 0.176 | 0.347 | 0.218 | 0.342 |
| 1.50 | 0.151 | 0.342 | 0.151 | 0.333 | 0.115 | 0.346 | 0.162 | 0.337 | 0.169 | 0.337 |
| 2.00 | 0.142 | 0.343 | 0.154 | 0.336 | 0.100 | 0.341 | 0.147 | 0.335 | 0.161 | 0.337 |

### 4 Supplementary figures

The main text emphasizes compact story-driven figures. To keep supplementary figures readable, dataset-specific panels are provided as separate figures rather than large multi-dataset composites.

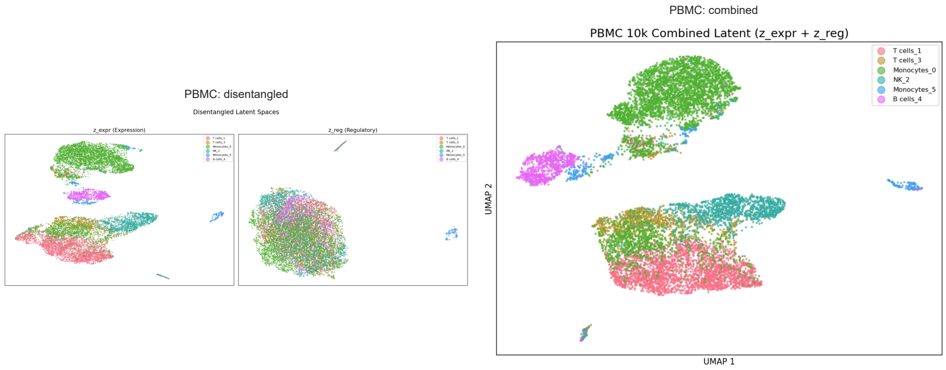


**Fig. S1** Dataset-specific UMAPs for PBMC showing the expression branch, regulatory branch and combined latent space.

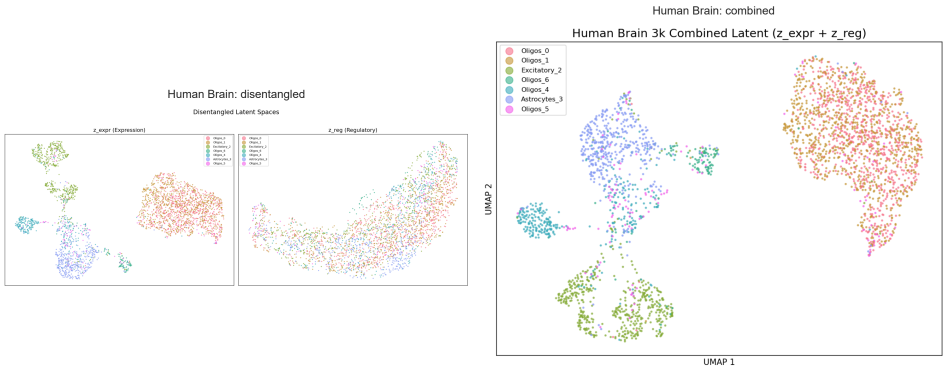


**Fig. S2** Dataset-specific UMAPs for Human Brain showing the expression branch, regulatory branch and combined latent space.

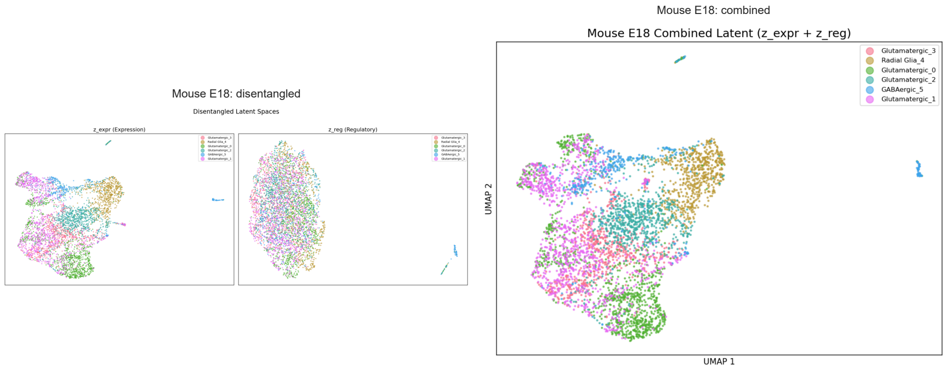


**Fig. S3** Dataset-specific UMAPs for Mouse E18 showing the expression branch, regulatory branch and combined latent space.

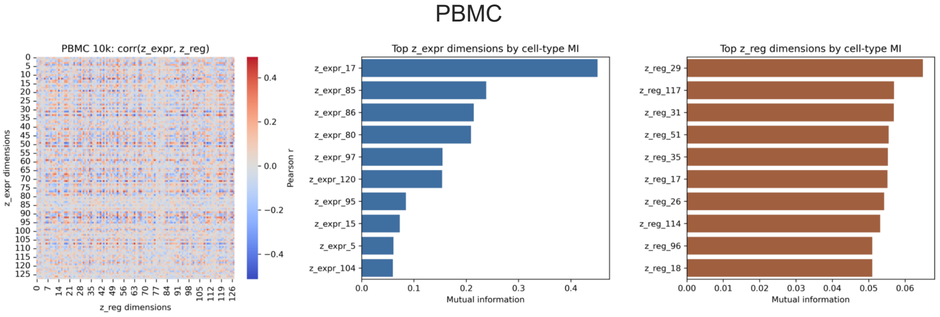


**Fig. S4** Full branch-separation analysis for PBMC, including cross-branch correlation and branch-specific benchmark-label association summaries.

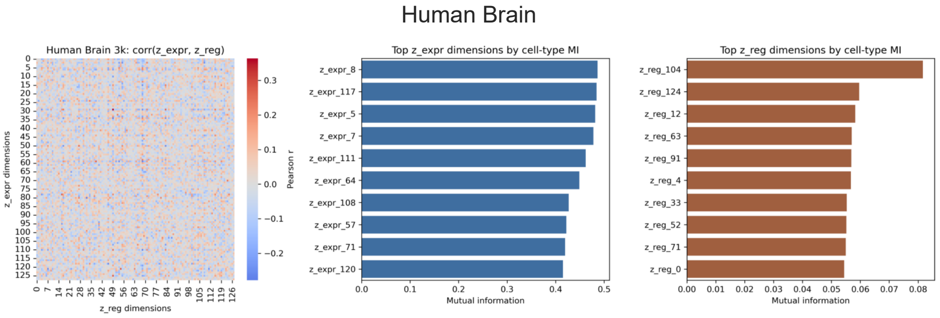


**Fig. S5** Full branch-separation analysis for Human Brain, including cross-branch correlation and branch-specific benchmark-label association summaries.

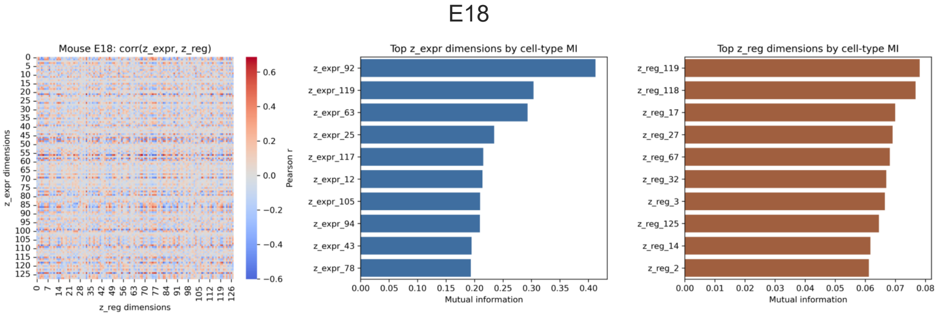


**Fig. S6** Full branch-separation analysis for Mouse E18, including cross-branch correlation and branch-specific benchmark-label association summaries.

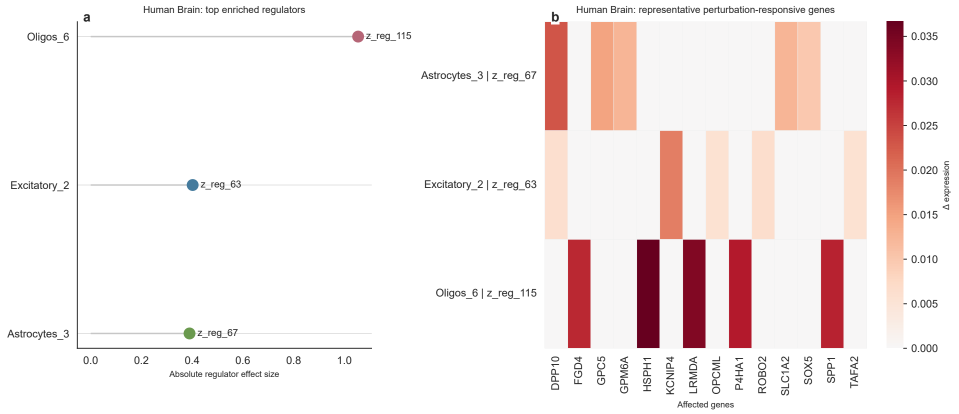


**Fig. S7** Human Brain discovery summary. The left panel shows representative cell-state-enriched regulators and the right panel shows representative gene-expression shifts after in silico knockdown, extending the regulator-centered analysis beyond PBMC.

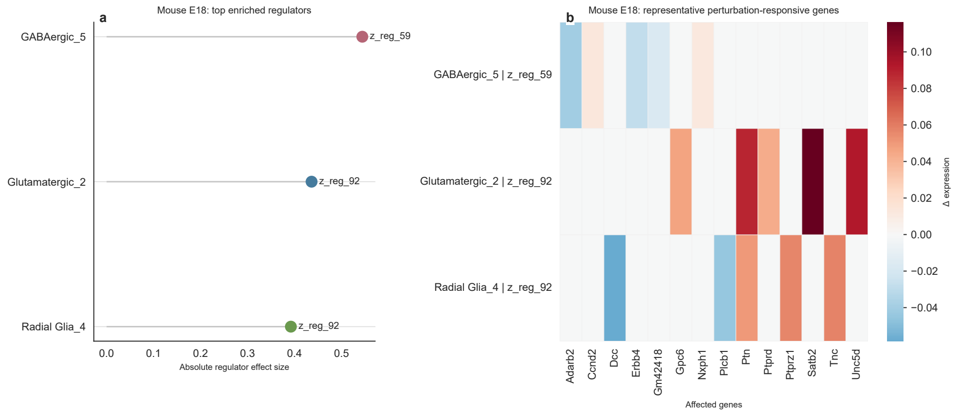


**Fig. S8** Mouse E18 discovery summary. The left panel shows representative cell-state-enriched regulators and the right panel shows representative gene-expression shifts after in silico knockdown, highlighting developmental-state programs in the regulatory branch.

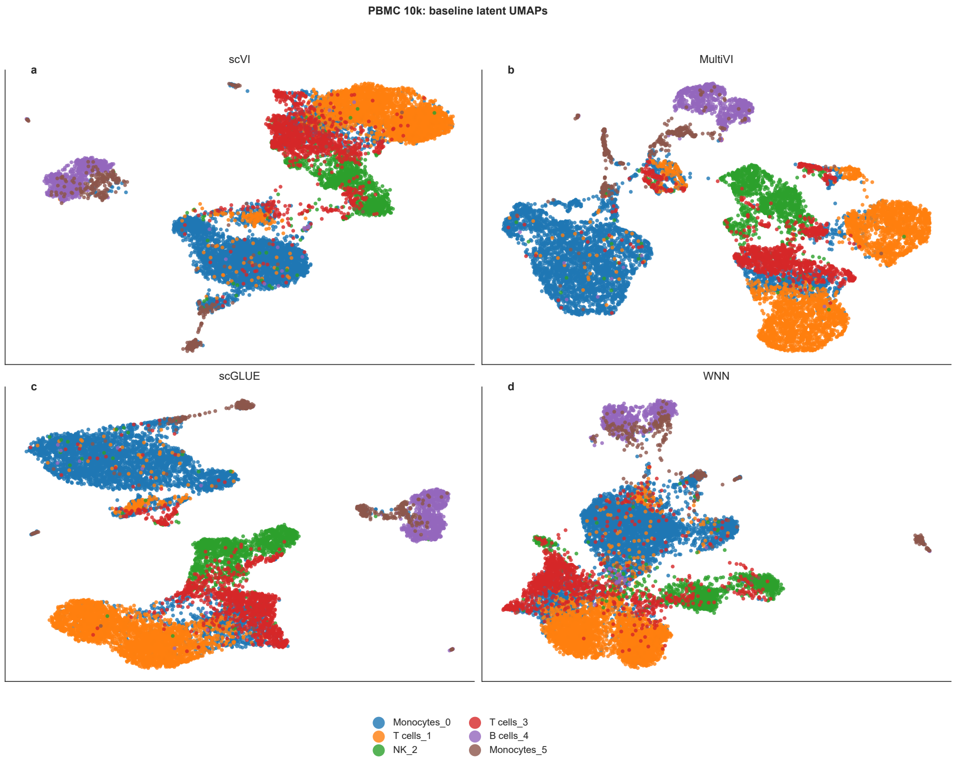


**Fig. S9** Baseline latent UMAPs for PBMC generated from scVI, MultiVI, scGLUE and WNN, shown as a visual reference for comparison with scDisent branch-specific embeddings.

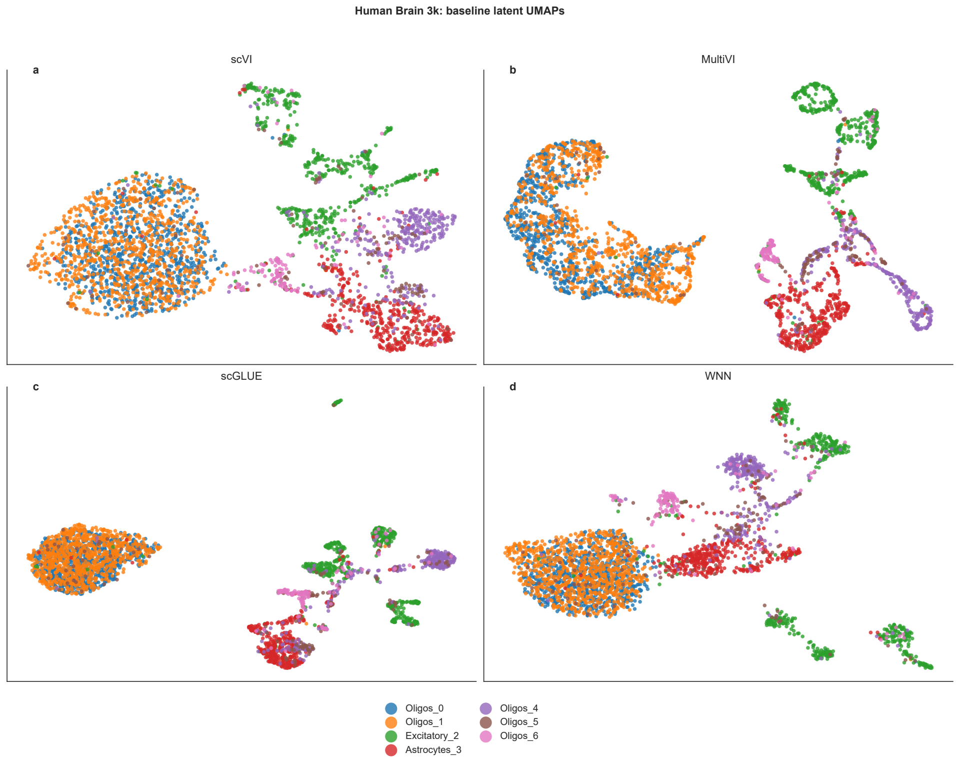


**Fig. S10** Baseline latent UMAPs for Human Brain generated from scVI, MultiVI, scGLUE and WNN, shown as a visual reference for comparison with scDisent branch-specific embeddings.

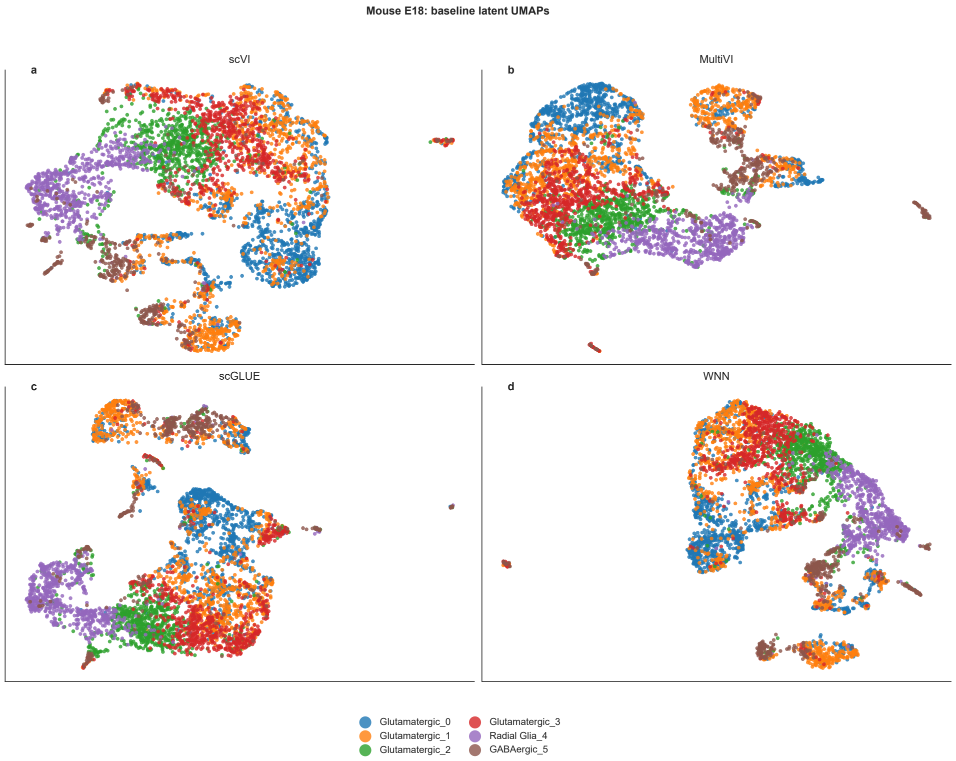


**Fig. S11** Baseline latent UMAPs for Mouse E18 generated from scVI, MultiVI, scGLUE and WNN, shown as a visual reference for comparison with scDisent branch-specific embeddings.

### 5 Supplementary notes on biological interpretation

The notes below provide a qualitative reading of the discovery outputs shown in the main text and supplementary figures. Their purpose is to interpret the regulator-centered summaries conservatively, connect them to verified marker biology when citations are available, and clarify where the study moves from benchmark evidence toward biologically motivated hypothesis generation. They are therefore interpretive notes rather than independent validation experiments.

#### Supplementary note 1. PBMC interpretation

The PBMC analyses are the strongest biological validation in this study because the relevant immune lineages are relatively mature and because the associated marker genes are well characterized. In this setting, the regulatory branch returns perturbation-oriented summaries that align with recognizable immune programs. The B-cell-associated regulator is linked to *BACH2*, *CD79A* and *MS4A1*; the NK-associated regulator is linked to cytotoxic markers such as *CCL5*, *NKG7* and *GZMH*; and monocyte-associated regulators connect to genes including *VCAN*, *LYZ* and *AOAH* (Muto et al. 2010; Vallejo et al. 2022; Chen et al. 2023). These are exactly the types of lineage-restricted summaries that the model is designed to expose.

For that reason, PBMC acts as the main reference case for asking whether the regulatory branch returns biologically coherent perturbation-oriented outputs. The answer appears to be yes: the highlighted latent factors are not random dimensions with weak marker overlap, but factors whose perturbation signatures are concentrated in lineage-relevant immune genes.

A separate resource-based cross-check against DoRothEA (Garcia-Alonso et al. 2019), summarized in `results/dorothea_validation_summary.txt`, found significant matches for all six PBMC regulators evaluated in that screen. Because the same best-matching TF recurred across those hits, we interpret this result conservatively as support that the perturbation signatures are structured and non-random, rather than as definitive regulator assignment.

These PBMC observations do not constitute causal proof, but they do support the claim that the regulatory branch is biologically organized rather than purely statistical. The main immune result is therefore not simply that scDisent can recover known markers, but that it can organize those markers into regulator-centered perturbation hypotheses rather than only clustering outputs.

#### Supplementary note 2. Human Brain interpretation

Human Brain extends that claim into a neural setting. Astrocyte-associated discovery outputs emphasize *GPC5*, *SLC1A2* and *SLC1A3*, while excitatory-neuron-associated outputs highlight genes such as *KCNIP4*, *ROBO2*, *DPP10* and *FGF12*. These patterns are concordant with atlas-level separation of astrocytic, excitatory-neuronal and oligodendroglial programs in the human brain (Chen et al. 2024). Multiple oligodendrocyte states also show distinct regulatory profiles rather than collapsing into a single shared signal. In particular, Oligos_6 yielded a large-magnitude regulator whose perturbation affected *HSPH1*, *LRMDA*, *P4HA1*, *SPP1* and *OLR1*, suggesting that scDisent is resolving state-specific neural programs within broadly related oligodendrocyte populations.

The brain setting is biologically important because it tests whether the same representational logic remains interpretable outside immune data. The relevant cell states are more heterogeneous and their boundaries are often softer than in PBMC (Chen et al. 2024). The fact that scDisent still returns astrocyte-, excitatory-neuron- and oligodendrocyte-associated regulators therefore argues that the method is not simply leveraging a particularly easy immune benchmark. Instead, it suggests that the regulatory branch can remain interpretable even when state structure is more continuous and atlas-like.

The main value of this section is therefore not that every neural regulator is definitively identified, but that the regulator-centered summaries stay coherent in a more atlas-like tissue. That is a stronger test of branch semantics than simply confirming that broad neural classes separate in a conventional manifold (Chen et al. 2024).

#### Supplementary note 3. Mouse E18 interpretation

Mouse E18 provides a developmental counterpart. A GABAergic-associated regulator was linked to shifts in *Adarb2*, *Erbb4* and *Foxp2*, whereas glutamatergic regulators highlighted genes such as *Satb2*, *Unc5d*, *Ptprd*, *Ptprk* and *Dcc*. Satb2 is a classic determinant of corticocortical projection-neuron identity (Leone et al. 2015), while *Tnc*, *Ptprz1* and *Fabp7* are compatible with radial-glia-like developmental programs (Pollen et al. 2015). The appearance of *Adarb2* in the GABAergic branch is also in line with inhibitory interneuron-associated populations (Machold et al. 2023). Together with the PBMC and brain results, these observations support the interpretation that scDisent is not merely learning cleaner clusters, but is exposing regulatory axes that can be queried in biologically distinct tissue contexts.

The developmental E18 setting is a useful stress test because transitional states are common and coarse reference labels do not fully capture the underlying biology (Pollen et al. 2015; Leone et al. 2015; Machold et al. 2023). In this context, the main value of scDisent is not that it perfectly resolves every developmental branch, but that it still returns regulator-centered summaries that can be related to neuronal lineage and progenitor programs. This is exactly the kind of setting in which a branch that is only weakly tied to coarse labels may be most informative, because it can encode state variation that is not exhausted by annotation.
